# The effect of pasture management on fire ant, *Solenopsis invicta* (Hymenoptera: Formicidae), abundance and the relationship to arthropod community diversity

**DOI:** 10.1101/2020.12.04.398214

**Authors:** Ryan B. Schmid, Jonathan G. Lundgren

## Abstract

The red imported fire ant, *Solenopsis invicta*, is one of the most prolific invasive species to the Southeastern U.S. These invaders preferentially colonize highly disturbed land and grassland habitat. Management of livestock in pasture systems can have a profound impact on the level of disturbance in grassland habitats, and we hypothesized that pasture management would have a significant effect on *S. invicta* abundance in the Southeastern U.S., and arthropod diversity would negatively correlate with this invasive species. We studied the effects that pasture management systems (based on stocking density, rotation frequency, and insecticide application rates) have on fire ant mound abundance and arthropod diversity for the soil, foliar, and dung communities. *S. invicta* mounds were quantified and mound areas were documented along transect lines in six pastures. Soil and foliar arthropod communities were collected along the same transect lines, and dung communities sampled from pats within the pasture system. Pastures managed under adaptive multi-paddock (AMP) practices had 3.35× more *S. invicta* mounds and 4.64× more mound area than their conventionally managed counterparts. However, arthropod diversity did not correlate with *S. invicta* abundance for any of the three arthropod communities sampled. This study shows pasture management can have a significant impact on *S. invicta* mound abundance, but arthropod communities in AMP managed pastures did not suffer decreased diversity from increased abundance of *S. invicta*. Additionally, this study demonstrates that this invasive species does not necessarily contribute to diversity decline, at least under AMP pasture systems.

## Introduction

Invasive species are regarded as one of the top threats to native biodiversity (Simberloff 2000). In general, the consensus is that invasive ants negatively affect the diversity of native ant communities and other arthropods (Porter and Savignano 1990, Human and Gordon 1997, Jourdan 1997, Hoffmann et al. 1999, Sanders et al. 2001). One of the most well-known invasive ants in the Southeastern U.S. is the red imported fire ant, *Solenopsis invicta* (Burden) (Hymenoptera: Formicidae), owing to environmental and economic problems associated with this species (Lofgren 1986). Since its invasion of the U.S. on the shores of Mobile, AL during the 1930s, this species has risen in notoriety from a relatively minor member of the South American ant community to become a dominant ant species throughout the Southeastern U.S. Many studies have found that the rise in population densities of *S. invicta* are negatively associated with overall ant species richness and abundance (Stein and Thorvilson 1989, Camilo and Phillips Jr. 1990). However, habitat disruption is a common confounding factor in these studies that can both increase *S. invicta* abundance while simultaneously hindering native ant communities (Wojcik 1983, Buczkowski and Richmond 2012). This is a difficult subject to detangle, as *S. invicta* is closely associated with disturbed habitats, and remains a poorly understood aspect of *S. invicta* ecology in the Southeastern U.S.

Open grassland environments are the preferred habitat of *S. invicta* in both the U.S. and its native range (Wojcik 1983). Consequently, livestock pastures represent a significant portion of *S. invicta*’s habitat in the Southeastern U.S. landscape. While *S. invicta* is typically described as a pest by many people in the U.S. (including livestock producers), this ant species can benefit certain aspects of cattle production on open pastures. First, reports from cattle ranchers indicated the appearance of *S. invicta* in pastures coincided with a decline in lone star tick populations. Subsequent studies found that foraging *S. invicta* significantly reduce lone star tick eggs, engorged larvae, and engorged female ticks in pastureland, which is likely the reason for the reduced lone star tick populations observed by ranchers (Harris and Burns 1972, Burns and Melancon 1977). Second, *S. invicta* reduces adult fly emergence of two major pest of the cattle industry, the horn-fly, *Haematobia irritans* (Linnaeus) (Diptera: Muscidae) and the stable-fly, *Stomoxys calcitrans* (Linnaeus) (Diptera: Muscidae) (Summerlin and Kunz 1978, Tschinkel 2006). Despite the impact of *S. invicta* on these dung-dwelling fly pests, *S. invicta* has not been shown to affect the reproduction and activity of beneficial dung beetles (Tschinkel 2006, Steele 2016). This suggests that *S. invicta* is beneficial for pest control in livestock production. While the impact of *S. invicta* on livestock operations has been studied, the reverse of this relationship or the degree to which the livestock management impacts the *S. invicta* community within pastures remains poorly understood. Given *S. invicta’s* affinity for disturbed habitats, it stands to reason that livestock management systems that frequently disturb pastures will increase the abundance of this invasive fire ant.

Adaptive multi-paddock (AMP) grazing is a grazing system that was developed and refined during the latter half of the 20^th^ century, due in part to mounting evidence for environmental benefits (Teague et al. 2011). AMP grazing utilizes herd management techniques like multiple paddocks per herd, high animal densities, short periods of grazing, adequate recovery periods for vegetation, and high stocking rates (Savory and Parsons 1980, Savory and Butterfield 1999, Teague 2014). AMP grazing practices improve the health of pasture soil and plant communities, which enhances soil organic matter, water holding capacity, nutrient availability, and increases fungal to bacterial ratios in the soil (Teague et al. 2011). These management practices also produce short, punctuated disturbances of paddocks within pastures, potentially creating the opportunity for *S. invicta* to seize upon the disturbed habitat and colonize the paddock following a grazing event. Understanding the impact of AMP grazing on the abundance of *S. invicta*, and consequently the impact of *S. invicta* on arthropod community diversity, is important to the sustainability of an AMP grazing system. Our hypothesis was that pastures managed under AMP grazing regimes would contain higher densities of *S. invicta* mounds relative to their conventional counterparts, and arthropod community diversity would negatively correlate with *S. invicita* mound abundance.

## Methods

### Site selection

Sampled pastures (n = 6) were located in two Southeastern U.S. states (Alabama n = 4, Mississippi n = 2) (Figure 1). Sampling occurred within the known range of *S. invicta* in the U.S., and *S. invicta* mounds were present in all sampled pastures. All grazing management systems implemented on sampled pastures were employed at least 10 years prior to sampling. Ranch managers executed a variety of grazing management practices that fit within their ranching system and goals. Management practices that varied between operations included stocking density, rotation frequency, and insecticide usage (Table 1). These management practices were used to categorize ranches into one of two treatment groups, AMP and conventional, based on meeting a majority (≥ 4) of the qualifications defined in Table 1. Grazing treatments were paired (< 8 km apart) between treatments across study locations.

**Table 1.**
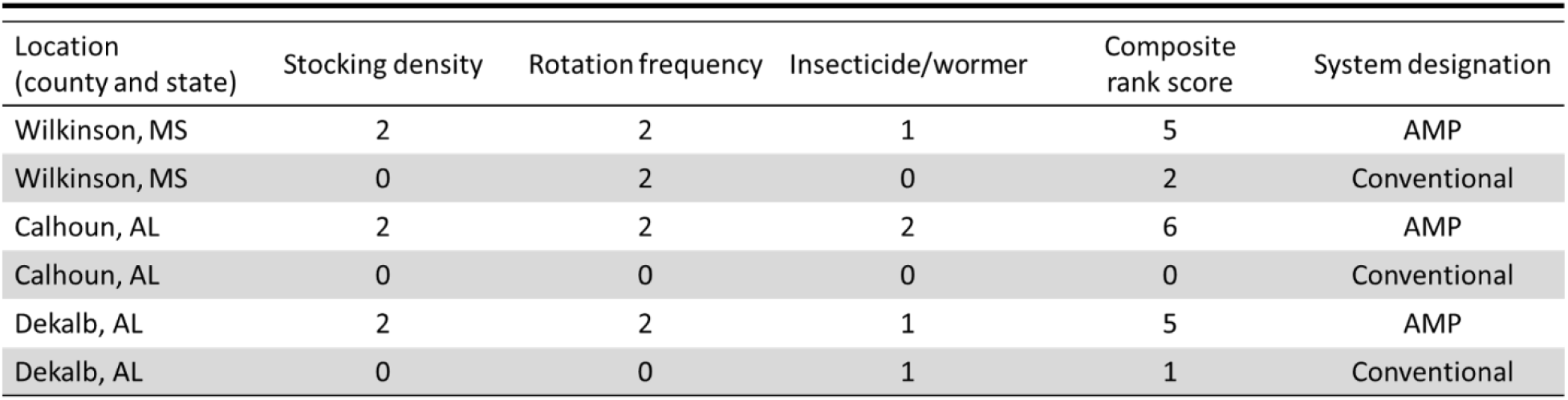
Ranch systems (n = 6) were categorized based on three cattle management practices: stocking density of cattle, rotation frequency of the herd, and use of insecticides to develop a composite rank score for each ranch. Ranches whose rank scores are in the top 50% were considered adaptive multi-paddock grazing (AMP) (white rows); those with rank scores in the lower 50% were considered conventional (shaded rows). The three management practices were scored 0 – 2, with higher numbers reflecting AMP practices. Stocking density (animal unit; AU) was divided into <5 AU/ha (0), 5 – 10 AU/ha (1), and >10 AU/ha (2). Rotation frequency was divided into >30 d rotation (0), 10 – 30 d rotation (1), and <10 d rotation (2). Insecticide/wormer application was divided into multiple applications (0), application once a year to individuals in herd thought to need treatment (1), and no insecticide or wormers (2).

**Figure 1.**
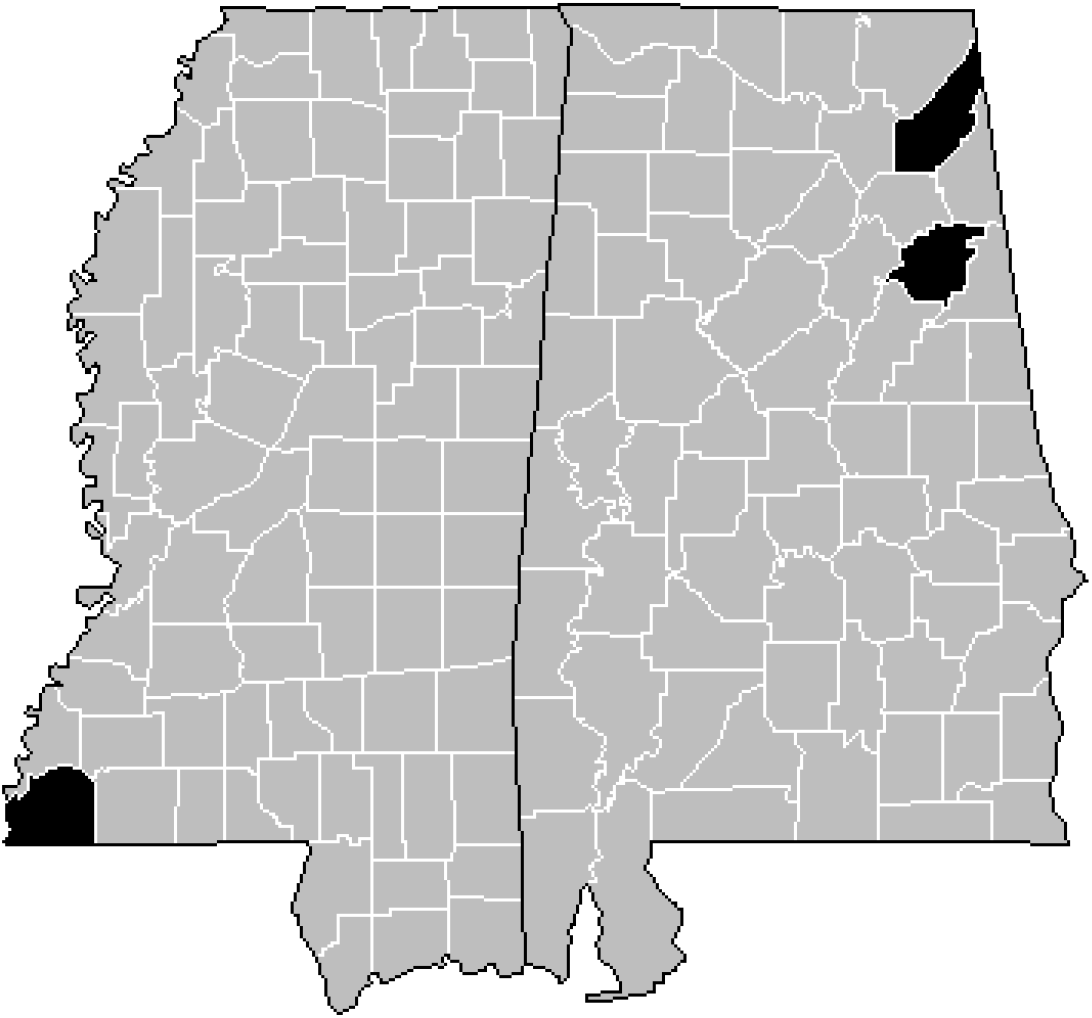
State maps of ranches sampled for *Solenopsis invicta* mound abundance and areas, as well as soil, foliar, and dung arthropod community diversity. Ranches were located within highlighted counties (Wikinson county, MS, n = 2 ranches; Calhoun county, AL, n = 2 ranches; DeKalb county, AL, n = 2 ranches).

### Sampling procedure

Sampling occurred from September 29 – October 1, 2018. Two separate areas with comparable soil characteristics within each pasture were selected for sampling. Three 46 m transect lines were established perpendicular to the slope of each of the two sampling areas, for a total of six transect lines sampled. *S. invicta* mounds present within 1 m of transect lines were counted, along with the area of each mound according to procedures outlined by Porter et al. (1992) for sampling fire ant mound densities and areas along transects. The soil arthropod community was sampled at 7.6 m and 38.1 m on four of the transects using soil cores (10 cm diameter, 10 cm deep). Dung arthropod community was sampled from randomly selected dung pats from each ranch (n = 5 pats per ranch). Age of the dung pats ranged from 2 – 5 d old, as this age of pat has peak arthropod abundance and diversity (Pecenka and Lundgren 2018). Arthropods within soil cores and dung pats were extracted using a Berlese system over 7 d, which ensured that each soil/dung core had completely dried and all arthropods had left the core. Soil and dung cores were kept cool on ice upon extraction from the field, and loaded into the Berlese system as soon as possible after extraction (within 60 h of collection). Foliar-dwelling arthropods were sampled from pasture vegetation at 22.9 m along three of the transect lines that *S. invicta* mounds were sampled. Pasture foliage was swept with a 38 cm diameter net, with 25 sweeps occurring perpendicular to each side of the transects (total of 50 sweeps per transect). Arthropods collected from sweep samples and extracted from the Berlese system were stored in 70% ethanol, until they could be identified and cataloged. Each specimen was identified to the lowest taxonomic level, representing functional morphospecies. Larvae were considered as distinct morphospecies, owing to their discrete differences in ecological function. Vouchers of all specimens are housed at the Mark F. Longfellow Biological Collection at Blue Dasher Farm (Estelline, SD).

### Data analysis

*S. invicta* mound abundance and area were compared between grazing treatments (AMP and conventional grazing) using one-way ANOVA, with the data conforming to the assumptions of ANOVA. Linear regression analysis was conducted to gain greater insight on correlations between *S. invicta* abundance and arthropod community diversity and species richness for soil, foliar, and dung arthropod communities. Statistical significance for *P*-value was set at α = 0.05. All statistics were conducted using Systat 13 (Systat Software, Inc.; Point Richmond, CA).

## Results

### Arthropod communities

The complete inventory of arthropod morphospecies collected from this study are presented in Schmid et al. (In prep). There were 15,705 arthropod specimens, representing 184 morphospecies, collected in the soil samples. In the foliar community, there were 13,376 arthropod specimens collected, representing 371 morphospecies. Lastly, dung arthropod sampling resulted in 3,465 specimens, representing 110 morphospecies. Acari and Collembola represented 14,983 and 3,603 specimens, respectively, of the total 32,546 collected arthropod specimens from the three communities. Lack of taxonomic keys and technical expertise to identify these two groups beyond the order Acarina and the family level for Collembola prevented the inclusion of these groups in diversity analysis.

Solenopsis invicta *mound abundance and area in livestock management systems. Solenopsis invicta* mound abundance differed significantly between grazing treatments (F_1,4_ = 11.77; *P* = 0.03) (Figure 2A). Mean ant mound abundance was 3.35× higher in AMP managed pastures than conventional pastures. The higher abundance of *S. invicta* mounds resulted in a significantly greater area of *S. invicta* mounds per pasture (F_1,4_ = 33.06; *P* = 0.01) (Figure 2B), with the mean area of *S. invicta* mounds measured within AMP pastures being 4.64× more than conventional pastures.

**Figure 2.**
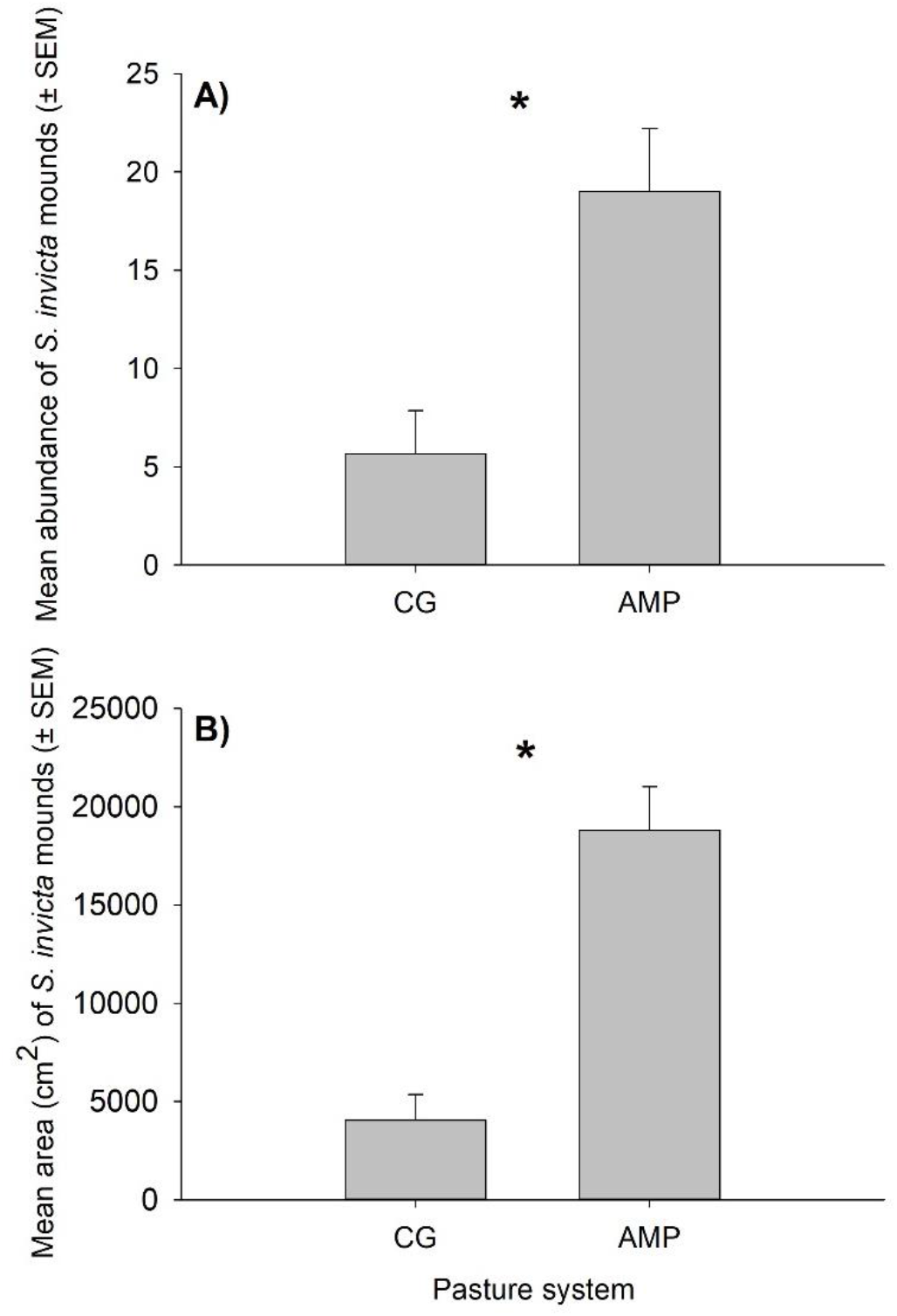
Mean (± SEM) *Solenopsis invicta* mound abundance (A) and mound area (B) in adaptive multi-paddock (AMP) and conventionally grazed (CG) pastures (n = 6). Statistical difference (*) was set at α = 0.05.

Solenopsis invicta *correlations to arthropod diversity*. Soil, foliar, and dung arthropod community species richness and diversity (Shannon H’) varied between sampled pastures. However, *S. invicta* mound abundance did not significantly correlate to species richness or diversity for any of the arthropod communities (Table 2).

**Table 2.** Regression analysis results comparing *Solenopsis invicta* mound abundance to arthropod community (soil, foliar, and dung communities) species richness and diversity (Shannon H’).

|  | <u>Soil Arthropods</u> |  |  | <u>Foliar Arthropods</u> |  |  | <u>Dung Arthropods</u> |  |  |
| --- | --- | --- | --- | --- | --- | --- | --- | --- | --- |
|  | df | F-ratio | P-value | df | F-ratio | P-value | df | F-ratio | P-value |
| Species Richness | 1, 4 | 0.04 | 0.86 | 1, 4 | 1.44 | 0.30 | 1, 4 | 1.24 | 0.33 |
| Diversity (Shannon H') | 1, 4 | 0.77 | 0.43 | 1, 4 | 1.47 | 0.29 | 1, 4 | 4.70 | 0.10 |

## Discussion

This study revealed significant differences in *S. invicta* mound abundance and area within pasture systems under differing cattle management in the Southeastern U.S. Higher mound abundance and area was consistently found in sampled areas within AMP managed pastures, relative to their conventionally managed counterparts. Despite higher abundance of *S. invicta*, none of the sampled arthropod communities (soil, foliar, and dung) correlated with *S. invicta* mound abundance. This result opposes our hypothesis that *S. invicta* abundance would negatively correlate with arthropod community diversity, which was founded upon the reputation of *S. invicta* to decrease native arthropod community diversity within invaded habitats (Camilo and Phillips Jr. 1990, Porter and Savignano 1990, Lu et al. 2012).

There are at least two potential explanations for the lack of correlation between *S. invicta* mound abundance and arthropod community diversity. First, *S. invicta* and arthropod communities may be governed by disparate regulatory factors in our tested pasture systems. For example, both biotic and abiotic factors, e.g., temperature, soil structure, moisture, habitat heterogeneity, are known to affect *S. invicta* and arthropod communities in myriad ways (Tschinkel 2006, Gurr et al. 2012). It is plausible that the range and complexity of environmental factors interacting within our pasture systems may affect *S. invicta* populations independently of the native arthropod communities. However, this hypothesis contrasts Morrison and Porter (2003), which documented positive correlations between *S. invicta* populations and the ant and non-ant arthropods, suggesting that *S. invicta* and other arthropods were regulated by common factors. It is worth noting that Morrison and Porter (2003) exclusively sampled on pasture land, and although grazing intensity varied among sites, grazing intensity was not included in the analysis of *S. invicta* nor arthropod diversity. The results of our study indicate that grazing intensity may be an important factor when examining *S. invicta* abundance in pasture land.

This leads us to hypothesize a second explanation for the lack of correlation between *S. invicta* and arthropod communities. We theorize the native arthropod communities measured in our AMP pasture systems may be more resilient to invading *S. invicta*, effectively able to buffer against invasion by a foreign species as a result of community or habitat structure. A keystone goal of AMP producers is to increase the resilience of an operation by bolstering various aspects of the environment, e.g., vegetation diversity, soil microbiota, soil physical properties, and water holding capacity (Teague et al. 2011, Hillenbrand et al. 2019). Strengthening these environmental features supports a more diverse array of flora and fauna (including arthropods) (Huston and Marland 2003, Welti et al. 2017, Aldebron et al. 2020), and this increased habitat complexity provides more niches, structure, resources, etc. that may have allowed the AMP arthropod community to maintain diversity while also absorbing higher numbers of *S. invicta*. As our study was based on correlation and not causality, we are unable to substantiate either theory with the data from this study.

Anecdotal evidence gathered via informal discussions with ranch managers at the time of sampling suggested that AMP managed pastures would have fewer *S. invicta* mounds, and consequently fewer associated problems with *S. invicta* in their pastures. Opinions on this subject were based upon experience and observations across the fence lines by the ranchers themselves, who observed *S. invicta* mounds around gates, water tanks, and well-trodden cattle-paths in their conventionally managed counterparts. Opposingly, AMP ranchers actively try to maintain ground cover throughout their operation through managed herd movement and elongated land rest periods; thus, minimizing high-traffic areas and bare ground (Gosnell et al. 2019). Often, these high-traffic areas are used by humans as well as livestock during routine checks on the herd, water tanks, mineral feeders, fencing, etc. The lack of abundant *S. invicta* mounds in areas frequented by AMP herdsmen may have led them to believe they had less fire ants in their pastures. However, our data shows the opposite, at least in our sampled areas away from high-traffic zones. The areas that were sampled in the pastures is an important point to keep in mind when interpreting our results together with the opinion of the ranchers. It may simply be that the AMP ranchers interviewed for this study do not come into contact with *S. invicta* in their pastures as often because mounds are not present in highly trafficked areas. Taking the opinion of ranchers into consideration with our results, we hypothesize that regeneratively managed ranches had greater functionality within their pastures because *S. invicta* did not hinder ranch operations with mounds present in high-traffic areas. This is an important hypothesis to consider for future study, as mitigating human interaction and *S. invicta* associated problems would be an important positive outcome of AMP ranching. While the functionality of pastures expressed by these ranchers was beyond the scope of this study, it is worth noting for further investigation in ranching systems.

## Acknowledgements

We thank Liz Adee, Mike Bredeson, Nicole Schultz, Alec Peterson, Sierra Stendahl, and Kassidy Weathers for collecting specimens. Dr. Kelton Welch identified the insect specimens for this study. Funding for this project came from the Carbon Nation Foundation.

## Data accessibility

Upon acceptance to Ecology and Evolution, data will be archived on Dryad and DOI will be included with this manuscript.

